# MDMD: a computational model for predicting drug-related microbes based on the aggregated metapaths from a heterogeneous network

**DOI:** 10.1101/2023.10.13.562158

**Authors:** Jiajie Xing, Xianguo Zhang, Juan Wang

## Abstract

Clinical studies have shown that microbes are closely related to the occurrence of diseases in the human body. It is beneficial for treating diseases by means of microbes to modulate the activity and toxicity of drugs. Therefore, it is significant in predicting associations between drugs and microbes. Recently, there are several computational models for addressing the issue. However, most of them only focus on drug-related microbes and neglect related diseases, which can lead to insufficient training. Here we introduce a new model (called MDMD) is proposed to predict drug-related microbes based on the Metapaths from a heterogeneous network constructed by using the data of Diseases, Microbes, Drugs, the associations of microbe-disease and disease-drug. The MDMD uses an aggregation of the metapath features that can effectively abundance the embedding of the features for different types of nodes and edges in the heterogeneous networks. Then, the MDMD uses the attention mechanism to mark the importance of the metapath vector for each node type which can improve the quality of feature embedding. Experimental results demonstrate that the MDMD improves accuracy by 1.9% compared with other models. The MDMD is also used to predict the microbes of two drugs Lamivudine and Tenofovir which are the antiretroviral drugs used to treat the Acquired Immune Deficiency Syndrome(AIDS). The results show that 90-95% of microbes are reported in the PubMed. Mycobacterium tuberculosis(Mtb) is a specific microbe only predicted by the MDMD. An online platform of the MDMD is available in https://mdmd2023.bit1024.top/, in which the source code of the MDMD and the data in the work can be downloaded.

**Author summary:** Microbes inhabit multiple organs of the human body that consist of bacteria, fungi, and viruses. Extensive research shows that the microbes can adjust the efficacy and toxicity of drugs to treat the disease. The efficient and accurate selection of drug-related microbes is important for drug research and disease treatment. However, screening of drug-related microbes relies on traditional lab experiments that are labor-intensive and costly. With the growth of high-throughput data, the research of drug-related microbes urgently needs a computational method in bioinformatics. However, most of them only focus on drug-related microbes and neglect related diseases, which can lead to insufficient training. Therefore, we propose a new method (called MDMD) based on the aggregation of the metapath to efficiently and accurately predict potential drug-related microbes within the microbes-disease-drug network.

## Introduction

Microbes include archaea, fungi, viruses, and protozoa [8]. Lots of research have shown that lots of microbes inhabit various organs of humans, such as the gastrointestinal, respiratory, and genitourinary tracts [1]. The composition and diversity of the microbiota can be influenced by multiple factors, such as dietary habits and medication. Microflora dysbiosis can increase the risk of diseases. For example, a high-sugar and high-fat diet can lead to microbiota dysbiosis in the gut and result in the diseases of obesity and diabetes. The microbes can affect the immune function of the human body, metabolic and digestive systems. The imbalance of the immune system can result in allergies and autoimmune diseases. The imbalance of the metabolic system can cause metabolic disorders, such as obesity and diabetes [5]. Therefore, they are parasitic and symbiotic relationships between microbes and humans, which have also been demonstrated in research on microbes and diseases [33]. Several studies have shown that the break of the symbiotic relationship also leads to other diseases, such as cardiovascular disease [2, 4, 47]), impaired bone development [6], and even the occurrence of cancer [7]. We know that drugs can efficiently treat these diseases. Recent studies have emphasized the crucial role of microflora in regulating drug activity and side effects [53]. So, it is a key issue to treat diseases by modulating the activity and toxicity of drugs in terms of microbes [54]. For example, Immune Checkpoint Inhibitors(ICIs) are drugs used to treat cancer. Studies have shown that microbes can modulate the effects of drugs through immune [3]. However, overuse of drugs can result in antimicrobial resistance. Therefore, it is fundamental to research the association between microbes and drugs for disease treatment.

Traditional methods are both high-cost and time-consuming for identifying the association between microbes and drugs based on biological experiments [55]. It is convenient to predict them using computational approaches. So far, lots of methods have been proposed to predict the drug-related microbes from microbe-drug networks. HMDAKATZ [9] uses the KATZ metrics of the nodes similarity from microbe-drug heterogeneous network to predict the associations between microbes and drugs. A supervised CATAPULT method proposed by [52] based on the KATZ to predict microRNA and disease associations. MicroRNA and disease associations can be computed by KATZ. However, unknown associations can learn features by the CATAPULT. A proposed method uses the metapath2vec [13] method to predict the association between microbes and drugs from a bipartite network. GCNMDA [12] incorporates a CRF (conditional random field) on a graph neural network (GNN) to aggregate representations of neighborhoods from the microbe-drug bipartite networks. EGATMDA [11] improves the GCNMDA using an attention mechanism to aggregate the node embedding from an input graph to optimize model performance. Graph2MDA [18] uses a self-encoder (VGAE) to improve the embedding of the outlier nodes in order to predict drug-related microbes. GSAMDA [10] constructs a heterogeneous network by computing the similarity of microbes and drugs based on the Hamming interaction profile (HIP) method. It uses the GAT autoencoder and the sparse autoencoder(SAE) to learn the features of the nodes and edges from the network. GACNNMDA [43] inspired by the GSAMDA uses two layers of GAT to learn features. NTSHMDA [16] integrates the topological similarity of the network by a random walk. DeepMNE uses kernel neighborhood similarity to construct disease similarity and lncRNA similarity networks based on the deep learning method [46]. The GNNs used by these works will disregard the types of different edges, which can result in the insufficient learning of multiple associations (such as microbe-disease-drug) in the heterogeneous networks.

Recently, lots of works extract the features of nodes and edges from the heterogeneous networks by means of metapaths to predict the associations between biological elements, such as the drug–target [34], the disease-miRNA interactions [35], and the lncRNA-disease [36, 48]. Metapath2vec [13] combines metapaths and word2vec to learn the features from the heterogeneous networks in order to capture the semantic relationships of words. HAN [14] uses a hierarchical attention network including node-level and semantic-level attention to learn the features from metapaths. These models only focus on the start node and the end node from the metapaths. They often neglect the features of intermediate nodes. Most of them only use one metapath and not consider the correlation between multiple metapaths.

Here we aim to improve the embedding features of nodes in the heterogeneous networks when predicting drug-related microbes. The MDMD first constructs the heterogeneous networks based on the microbes, diseases and drugs, and maps the three types of nodes with different dimensions (i.e., microbes, diseases and drugs) into the same vector space. The MDMD then uses an encoder of metapath instances to learn the embedding features of intermediate nodes. It uses a multi-headed attention mechanism to calculate the attention weights among metapaths. The attention mechanism can improve the information loss between the metapaths. Finally, it uses a logarithmic loss to measure the difference between the predicted and true values. The result is a non-negative real number to indicate the degree of classification accuracy. We performed 5-fold cross-validation on MDAD [15] and aBiofilm [44] datasets.

## Materials

### Heterogeneous Networks

A heterogeneous network consists of different types of nodes and edges, denoted as *𝒢*= (*V, E*). where *V* is its node set and *E* is its edge set. The type of nodes is represented as a mapping *ϕ* from *V* to a set of node types *V T* . The type of edges is represented as a mapping *Ψ* from *E* to an edge type set *ET* . It is important to note that |*V T*| + |*ET*| *>* 2. It ensures at least two different types of nodes or edges in the heterogeneous network.

### Metapaths

Given a heterogeneous network *𝒢*, a metapath of *𝒢* is defined as 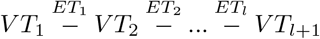 [17], *V T*_*i*_ (1 *≤ i ≤ l* + 1) are the type of nodes, and *ET*_*i*_ (1 *≤ i ≤ l*) are the type of edges. The pair of target nodes (*V T*_1_, *V T*_*l*+1_) can be represented by *ET* = *ET*_1_ *° ET*_2_ *°* …; *° ET*_*l*_ with the length *l* of the metapath, where denotes the operator between the edges.

### Metapath Instances

A metapath instance *R* is a path that connects multiple nodes in the heterogeneous networks, which describes the relationship including multiple types of nodes and edges.

## Method

### Data

We collect microbes-disease associations from MicroPhenoDB [19], HMDAD [20] and Disbiome [21], and obtain 5530 associations between 1773 microbes and 499 diseases. The names and treeID of 499 diseases are collected from the MeSH database [22]. We collect 9384 drugs-disease associations between 247 drugs and 127 diseases from the Drugbank [23], the CTD [24] and the Drugvirus.info [32]. There are 247 drugs and the 5586 drug-drug associations from the Drugbank and the ChEMBL [27]. We download the protein interactions and gene families of the related microbes from STRING database [49]. The detailed numbers of data are illustrated in Table 1.

**Table 1.** The detailed numbers of all data used by the work.

| Networks | Microbe | Disease | Drug | associations |
| --- | --- | --- | --- | --- |
| Microbe-disease | 1773 | 499 | - | 5530 |
| Microbe-microbe | 1773 | - | - | 1580 |
| Disease-disease | - | 499 | - | 3744 |
| Disease-drug | - | 127 | 247 | 9384 |
| Drug-drug | - | - | 247 | 5586 |

### Construction of the Heterogeneous network

#### Similarity of microbes

We use the method [25] to calculate the functional similarity among microbes based on their protein interactions and gene families. The genes of microbes may belong to different gene families. They have been encoded through the genome. There are interactions of genes in a gene family. The similarity of two microbes *M*_1_ and *M*_2_ is obtained by the equation 1.

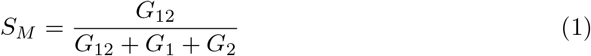

where *G*_12_ is the number of protein interactions encoded by gene families from the two microbes. *G*_1_ (*G*_2_) is the number of protein interactions encoded by gene families only from the *M*_1_ (*M*_2_).

#### Similarity of diseases

The method proposed by [45] is used to calculate the similarity of diseases. We first collected the synonym of diseases from Mesh databases that are tree structures. The disease is a node in the tree, while the synonyms are child nodes. The similarity *S*_*w*_ of the two diseases *v*_*i*_ and *v*_*j*_ can compute as follow:

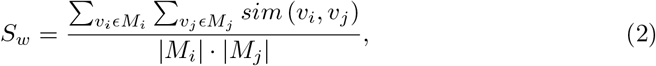

where *M* represents the number of synonyms for disease *v*. We compute the similarity between all possible diseases pair.

#### Similarity of drugs

SIMCOMP2 [37] is calculated the structural similarity for drugs, denoted as *S*_*Ds*_. *DIP* (Drug Interaction Profile) [51] is a vector of molecular fingerprints of drugs that are used to compute the Euclidean distance between the drugs *d*_*i*_ and *d*_*j*_. *S*_*Dg*_ is the Gaussian kernel similarity as follows.

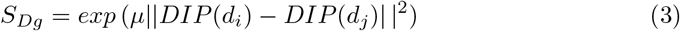

where *μ* is the width of the Gaussian kernel and is set as 1. The similarity of the two drugs is computed as the mean of *S*_*Ds*_ and *S*_*Dg*_.

### MDMD

Figure 2 shows the MDMD framework. It constructs a heterogeneous network, i.e. a Microbe-Disease-Drug network (denoted as M-Di-D). The layer of microbes in the network is based on the functional similarity of microbes, i.e. an edge between two microbes if their functional similarity is more than 0.5. The layer of diseases (drugs) is based on their similarity of diseases (drugs), i.e. an edge between two diseases (drugs) if their similarity is more than 0. The layers of between microbes and diseases, between diseases and drugs are constructed by the data described by the Data section, i.e. an edge between two different types of nodes if they have an association. The MDMD maps the different dimensions of features into the same dimension of features for three types of nodes in the M-Di-D network. It adopts the metapaths instance encoders to extract node content and edge relationships from the metapaths. It also uses an attention mechanism to aggregate the relationships of the various metapaths.

#### Feature mapping

The similarity values of microbes, diseases, and drugs are used as features of those nodes, which leads to different types of nodes having different dimensions of features. The MDMD uses the method proposed by Glorot [50] to map the different dimensions of features into the same dimension of features. The method uses a weight matrix which one value is from a normal distribution and an activation function to adjust the result dimension. Then all nodes have the same dimension of features 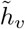.

#### Features of the metapaths

Figure 1 shows all metapaths in the M-Di-D network. Each metapath describes the associations of different node types. We design six metapaths that show the topology structure and potential features of the heterogeneous networks. *R*(*v, u*) denotes a instance of the metapath *R*, which describes the connection between the target node *v* and its begin node *u. R*(*v, u*) = (*v* = *l*_1_, *l*_2_ *…, l*_*m−*1_, *u* = *l*_*m*_). *m* is the number of nodes. *ET*_*i*_ is the edge between the nodes *l*_*i−*1_ and *l*_*i*_. *L*(*v, u*) is a sequence of nodes and edges without the first and the last nodes (i.e., v,u) of *R*(*v, u*). *et*_*i*_ is the type vector of edges related by the node *l*_2_. RotatE [26] is an encoder that can rotate the vectors of nodes and metapaths into a complex plane. 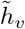 denotes the the mapping vector of the node *v*. Θ_*i*_ is the feature vector computed by the RotatE for the node *l*_*i*_ as follows:

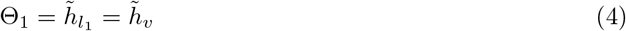

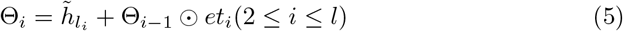

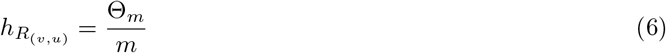

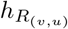 is the feature vector of the 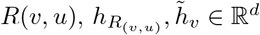. *⊙* the operation of multiplying for one to one component of two vectors. The GAT (Graph Attention Network) was proposed by [31] based on an attention mechanism. It indicates the importance of the matepath instance *R*(*v, u*) by assigning a weight vector 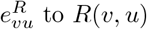 to *R*(*v, u*).So, the MDMD uses the attention mechanism of the GAT to assign a weight vector for *R*(*v, u*).

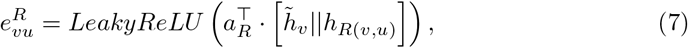

*a*_*R*_ *∈* ℝ^2*d*^ denotes the attention vector. The operator || is used to splice the vector 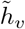 of the target node and the vector *h*_*R*(*v*,*u*)_ of the *R*(*v, u*). A softmax function is used to normalize the weight 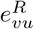 as 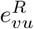 . Then the activation function *σ*(*·*) is used to aggregate the feature vectors of the metapath instances for the node *v*. Here we obtain the vector representation of the node *v* based on all metapath instances.

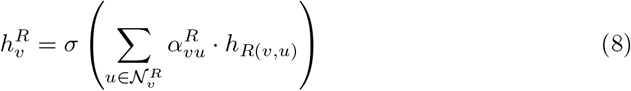

where 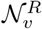 is the set of the nodes that are connected to the node *v*. Multi-head attention can stabilize the learning process. We extend the attention of equation 8 to multi-attention. The k-head attention is the following:

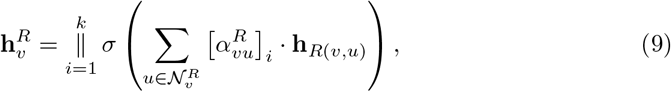

where *k* is a hyperparameter. For nodes *v*_1_, *v*_2_, …, *v*_*n*_ of the same type *i*, we compute a vector of metapath instances for each node and obtain a metapath vector *Z* for the *i*, i.e. 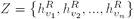.

**Fig 1.**
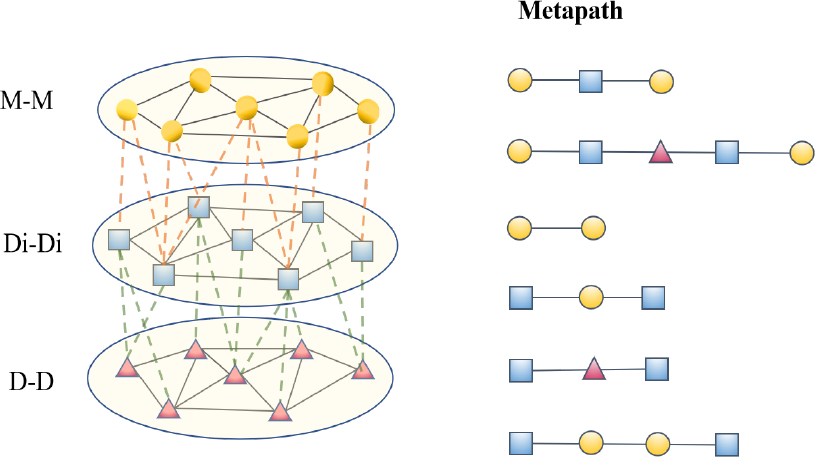
All metapaths in the heterogeneous network.

**Fig 2.**
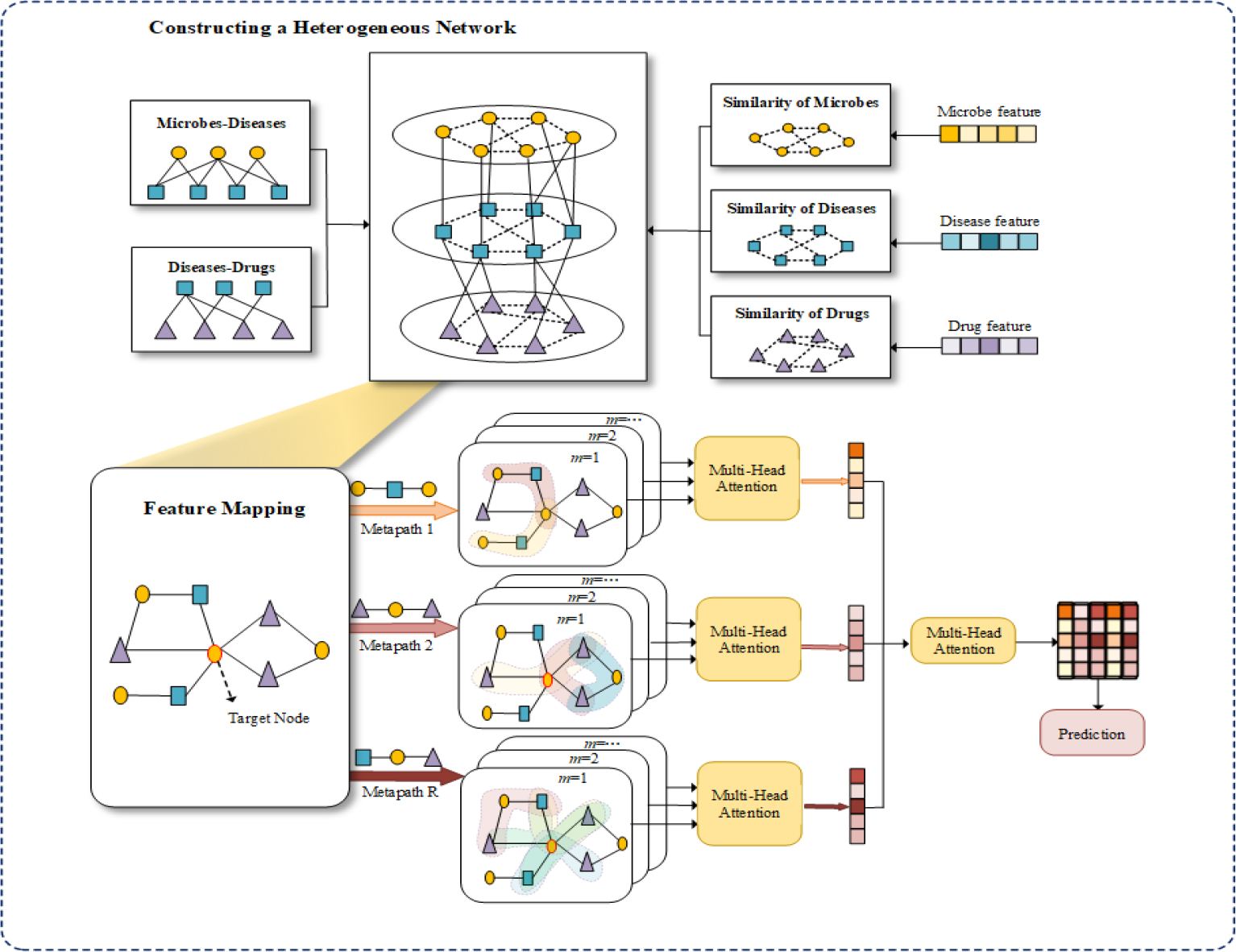
The framework of MDMD. First, we construct a heterogeneous network by the feature similarity of microbes, diseases, and drugs. Second, we map the different dimensions of features into the same dimension of features. Then, the MDMD extracts the features of the metapaths based on the metapath and attention for aggregation of the features between metapaths. Last, we obtain the embedding of the features to predict the drug-related microbes.

#### Aggregation of the features between metapaths

For a node type *i*, a non-linear activation function tanh is used to compute an average vector **S**_*i*_ of *Z* for *i*.

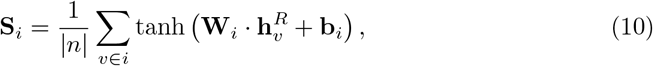

where 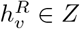. *W*_*i*_ and *b*_*i*_ are learned parameters, here *W*_*i*_ is a weight matrix and *b*_*i*_ is a bias term. *n* represents the number of nodes in *i*.

The attention vector of the learned parameters for the node type *i*, denoted as *q*_*i*_. *e*_*i*_ describes the importance for the vector *S*_*i*_, that can be computed by the similarity value between *q*_*i*_ with *S*_*i*_ as follow.

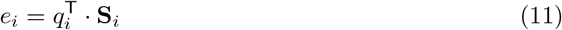

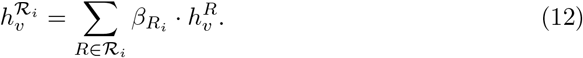

where 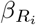 is a value normalized by a softmax function for *e*_*i*_. We assign weights to the each metapath *R*_*i*_ and summation. Then, we used a nonlinear activation function *σ*(*·*) mapping the vector 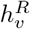 to the vector space as follow:

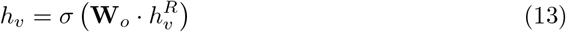

where *W*_*o*_ is a parameters matrix. The embedding vectors *h*_*v*_ is used to predict the drug-related microbes. Given a microbe *m* and a drug *d*, The predicted score *s*_*md*_ is calculated as:

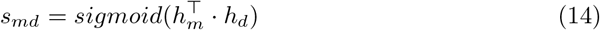

where *h*_*m*_ (*h*_*d*_) is the embedding of vector for *m* (*d*). We utilized the logarithmic loss function to enhance the accuracy of classification. A positive sample is an existent association between the node pairs(drug and the microbe), denoted as 1. The negative sample is not an existent association between the node pairs, denoted as 0.

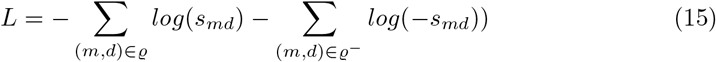

*ϱ* and *ϱ−* are the positive sample set and the negative sample set, respectively.

## Experimental

### Experimental design

We use 5-fold and 10-fold cross-validation (cv) on the MDAD and the aBiofilm datasets to evaluate the performance of models and reduce evaluation bias caused by uneven dataset partitioning. 10-fold cv is that the dataset is divided into 10 subsets, where the nine subsets are used to train the model and the remaining one is used to test the model. Two indexes AUC and AUPR are used to evaluate the performance of the MDMD. AUC is the area under the ROC curve, with values ranging from 0.5 to 1. The closer the value is to 1, the better the performance of the model. AUPR is the area under the Precision-Recall curve and is suitable for handling imbalanced datasets.

### Selection of hyperparameters

Hyperparameters control the performance of the model. Here we focus on three primary hyperparameters including the learning rate, dropout rate, and the number of attention heads. The dropout rate can lead to overfitting or underfitting issues if it is too high or too low, respectively. The number of attention heads with an excessively high value may lead to overfitting or high computational complexity. An experiment of 10-fold cv is done on the MDAD to assess the influence of these hyperparameters. The detailed experimentation results are shown in Figure 3. The AUC value is the highest when the learning rate is 0.0001. It indicates a slow learning process that can learn more features. A more significant step length may cross the optimal parameter or oscillate nearby the optimal local value. The optimum rate of dropout value is 0.5. The number of attention heads is 8.

**Fig 3.**
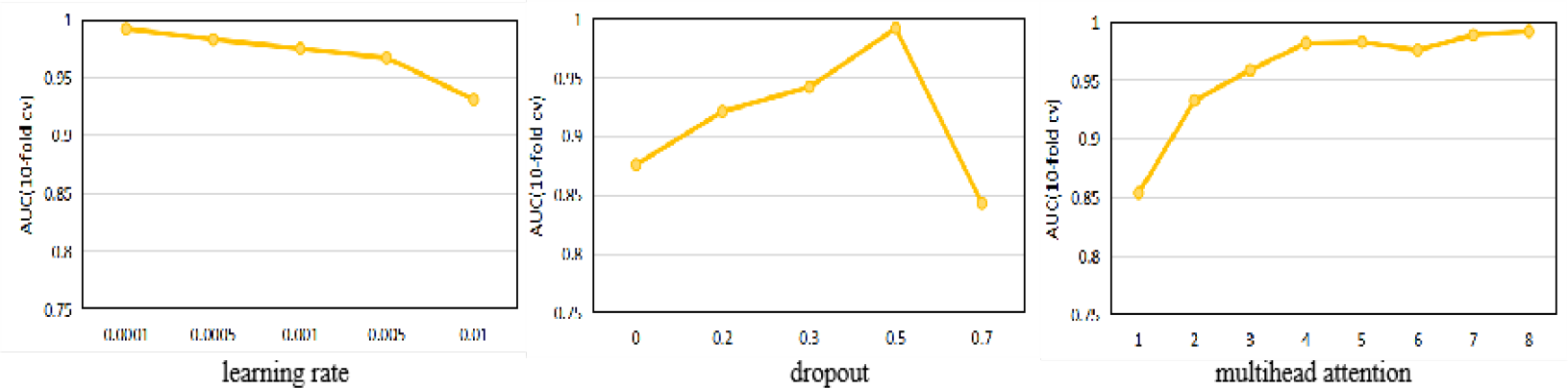
The impact of hyperparameters on performance.

### Performances of comparison experiments

The MDMD is compared with six state-of-the-art computational methods on the aBiofilm and MDAD databases, including the Graph2MDA, the GCNMDA, the EGATMDA, the GSAMDA, the HMDAKATZ, and the NTSHMDA. Default parameters are used to ensure the fairness of the experimental results. The experiments use a 5-fold cv. Table 2 displays the results of the experiments. Among all methods, the MDMD achieves the best performance on the aBiofilm dataset with AUC 0.9893±0.0191 and AUPR 0.9847±0.0106, which are 2.91% and 1.78% higher than the second-best method, Graph2MDA. The results of the MDAD dataset show that the MDMD with AUC 0.9923±0.0071 and AUPR 0.9942±0.0136, which are 1.91% and 1.43% higher than the second-best method.

**Table 2.** The results of the AUC and AUPR values for aBiofilm and MDAD datasets on comparison.

|  | aBiofilm |  | MDAD |  |
| --- | --- | --- | --- | --- |
|  | AUC | AUPR | AUC | AUPR |
| **MDMD** | 0.9893±0.0191 | 0.9847±0.0106 | 0.9923±0.0071 | 0.9942±0.0136 |
| Graph2MDA | 0.9602±0.0016 | 0.9799±0.0002 | 0.9732±0.0011 | 0.9764±0.0137 |
| GCNMDA | 0.9425±0.0412 | 0.9213±0.0018 | 0.9423±0.0131 | 0.9341±0.0071 |
| EGATMDA | 0.9643±0.0092 | 0.9631±0.0621 | 0.9605±0.0023 | 0.9535±0.0009 |
| GSAMDA | 0.9581±0.0681 | 0.9661±0.0537 | 0.9643±0.0038 | 0.9563±0.0011 |
| HMDAKATZ | 0.9104±0.0007 | 0.9183±0.0041 | 0.9283±0.0004 | 0.9232±0.0432 |
| NTSHMDA | 0.6744±0.0082 | 0.6798±0.0312 | 0.6897±0.0210 | 0.6741±0.0038 |

### Ablation analysis

Here we analyze performs of extracting features from each metapath (F), encoding the metapaths (E), and feature aggregation between metapaths (A). The MDAD used a 10-fold cv on the MDAD dataset. Figure 4 shows the results of the MDMD and its variants. The MDMD-F does not extract features from each metapath that showed poor performance compared to the MDMD. The MDMD-EM uses a mean encoder. The MDMD-EL uses a linear encoder, and the MDMD-ER uses a RotatE encoder of relational rotation. Among the three encoders tested, the MDMD-ER achieved the highest performance by preserving the structure of the metapath. The linear encoder(MDMD-EL) outperformed the mean encoder(MDMD-EM) by adding a linear transformation to the mean encoder. The MDMD-A indicated that feature aggregation between metapaths improves the performance of the MDMD. The success of the MDMD can be attributed to feature extraction from metapaths, encoding of the metapaths by the RotatE, and feature aggregation between the metapath.

**Fig 4.**
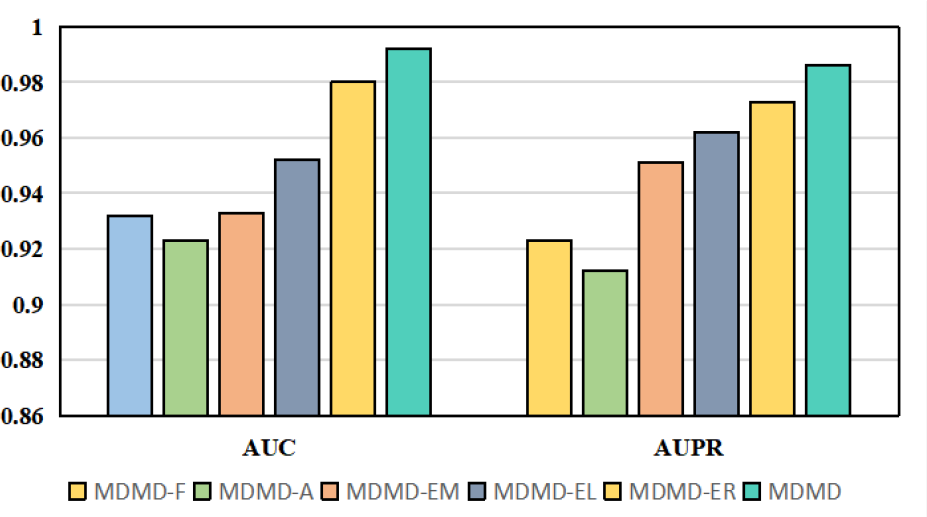
The results on the AUC and the AUPR values for ablation experiments.

### Case analysis

The MDMD is used to further validate the effectiveness in predicting drug-related microbes. First, we predict the antibacterial drugs Oxacillin and two drugs Lamivudine and Tenofovir to treat the disease of HIV. Second, we search related literature predicted by the MDMD for the top 10 and 20 microbes from the PubMed database. We then record the PMID numbers of the literature. The results are shown in Table 3, and the results of the HIV drugs are shown in Table 4. Existing research has shown that Oxacillin is a *β*-lactam antibiotic commonly used to treat bacterial infections, such as streptococci, staphylococci, and E. coli [28]. Oxacillin works by inhibiting the cell wall synthesis of bacteria to work through its medicinal effects [29]. Bacteroides fragilis is an enteric bacterium that can cause abdominal infections. Combined with other antibiotic drugs, it can treat bacterial infections [30]. The disease of Acquired Immunodeficiency Syndrome (AIDS) is caused by the human immunodeficiency virus (HIV), which is a viral infection [39]. HIV causes the immune system to lose its function and makes the patient susceptible to diseases [41]. HIV is an RNA virus that transcribes RNA into DNA and inserts it into the genome of the host cell, so the HIV virus is difficult to completely remove [42]. Recently, many drugs, including antiretroviral therapy (ART) such as Lamivudine and Tenofovir, have been used to relieve symptoms of the disease by effectively inhibiting the replication and multiplication of HIV [38]. Patients with AIDS often experience infections caused by Candida and Pneumocystis pneumonia infection [40].

**Table 3.** 20 microbes associated by Oxacillin.

| Microbes | PMID | Microbes | PMID |
| --- | --- | --- | --- |
| Bacteroides | 6142862 | Escherichia coli | 34870473 |
| Bacteroides fragilis | 2759932 | Firmicutes | NA |
| Burkholderia cepacia | 28634144 | Haemophilus influenzae | 34550809 |
| Citrobacter freundii | 36216208 | Klebsiella pneumoniae | 31010862 |
| Clostridium perfringens | 11324682 | Eggerthella lenta | NA |
| Porphyromonas gingivalis | 10412852 | Pseudomonas putida | 12919763 |
| Eikenella corrodens | 16962664 | Staphylococcus aureus | 28893360 |
| Staphylococcus epidermidis | 31462553 | Enterobacter aerogenes | 12462424 |
| Enterococcus faecalis | 31759592 | Streptococcus mitis | 9537382 |
| Enterococcus faecium | 31718570 | Streptococcus mutans | 24761805 |

**Table 4.** The related microbes of Lamivudine and Tenofovir.

| Lamivudine | PMID | Tenofovir | PMID |
| --- | --- | --- | --- |
| Escherichia coli | 2441708 | Streptococcus pneumoniae | 21471820 |
| Firmicutes | 10879645 | Klebsiella oxytoca | 31760254 |
| Shigella dysenteriae | 24417087 | Bacillus atrophaeus spore | 22226641 |
| Klebsiella pneumoniae | 19526705 | Bacteroides spp | 28732773 |
| Mycobacterium tuberculosis | NA | Firmicutes | 10879645 |
| Staphylococcus aureus | 19526705 | Gardnerella vaginalis | 28732773 |
| Staphylococcus epidermidis | NA | Streptococcus agalactiae | 30315206 |
| Streptococcus pneumoniae | 10585995 | Streptococcus oralis | 30315206 |
| Bifidobacterium adolescentis | 24417087 | Streptococcus mitis | 30315206 |
| Shigella dysenteriae | 24417087 | pathogens cytomegalovirus | 21471820 |
| Pseudomonas aeruginosa | 19526705 | Neisseria meningitides | 21471820 |

## Conclusion

In this study, we propose a new method called the MDMD, which is proposed to predict drug-related microbes using a heterogeneous network constructed by diseases, microbes, drugs, the associations of microbe-disease and disease-drug. The MDMD enriches the features of the microbes-disease-drugs heterogeneous network. The aggregated of the metapaths can effectively abundance the embedding of the features for different types of nodes and edges in the heterogeneous networks. The attention mechanism to mark the importance of the metapath vector for each node type can improve the quality of feature embedding. The experimental results show that the MDMD accurately predicts drug-related microbes, achieving an average of 99.2% improves accuracy by 1.9% compared with other models. The success of the MDMD can be attributed to feature extraction from metapaths, encoding of the metapaths by the RotatE, and feature aggregation between the metapath in the ablation analysis. In the case analysis, the MDMD is used to predict the microbes of two drugs Lamivudine and Tenofovir which are antiretroviral drugs used to treat the Acquired Immune Deficiency Syndrome(AIDS). The experimental results showed that 90-95% of the drug-related microbes are reported in the PubMed database. Interestingly, the Mycobacterium tuberculosis(Mtb) is a specific microbe only predicted by the MDMD, as Fig. 5. The antiretroviral drugs are one of the effective ways to treat the disease of pulmonary tuberculosis(PTB) caused by the Mtb [56]. The Lamivudine is an antiretroviral drug that is used to treat AIDS. We infer that the Lamivudine may be used to treat the PTB caused by the Mtb. This finding provides further evidence that the MDMD used features of disease helps to explore the potential relationship between disease-microbe, and disease-drug.

**Fig 5.**
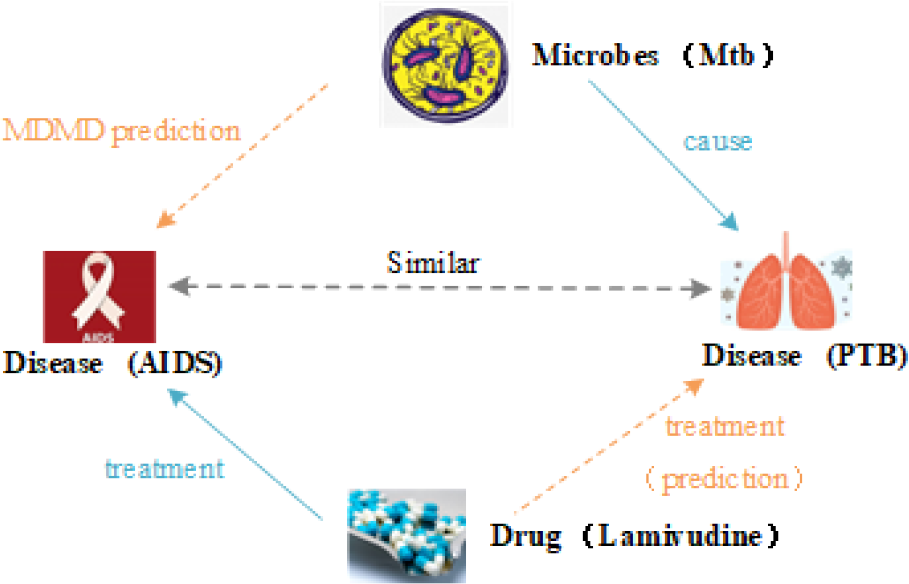
The examples of case analysis. The MDMD found that the Mtb causes the diseases of FTB and AIDS. The Mtb was associated with the drug Lamivudine in the results of the MDMD prediction. The Lamivudine is always used to treat the AIDS.

## An online platform for predicting

We designed an online platform of MDMD used to predict drug-related microbes is available in https://mdmd2023.bit1024.top/. The online platform accepts the chemical structure of drugs as input, which can be obtained from databases such as ChEMBL, and DrugBank. The source code of the MDMD and the data in the work can be downloaded which provides a convenient tool for relevant research.

## Acknowledgments

This work was supported by the National Natural Science Foundation of China (62002181 and 62061035).

